# Using cryogenic electron microscopy methods to gain insight into structure and initial host attachment of the flagellotropic bacteriophage 7-7-1

**DOI:** 10.1101/2025.01.29.635402

**Authors:** W.E.M. Noteborn, R. Ouyang, T. Hoeksma, A. Sidi Mabrouk, N.C. Esteves, D.M. Pelt, B.E. Scharf, A. Briegel

## Abstract

Understanding the structural and functional mechanisms of bacteriophage 7-7-1, the flagellotropic phage infecting *Agrobacterium* sp. H13-3, offers promising insights into phage-host interactions. Using single particle analysis (SPA) cryo-electron microscopy (cryo-EM), we determined the capsid and tail structure, and built atomic models of capsid hexamers, pentamers and tail. Combined with cryo-electron tomography (cryo-ET) and machine learning methodologies, our findings indicate that phage 7-7-1 uses capsid fibers to establish initial contact with the host flagellum, followed by subsequent attachment to cell surface receptors. Proteinase K treatment confirmed the time-dependent degradation of capsid fibers. The study also demonstrated that capsid fibers are flexible and can interact with other phages and host flagella, suggesting a cooperative infection strategy. These results provide crucial structural insights and may open avenues for developing phage-based therapeutics against resistant bacterial pathogens.

## Introduction

Flagellotropic bacteriophages represent a distinct group of phages that initiate their infection cycle by attaching to the flagellum of their motile host. The bacterial flagellum, a helical extracellular appendage powered by a rotary motor used for swimming and swarming, serves as the initial point of attachment for these phages. Exploiting the rotational motion of the flagellum, flagellotropic phages navigate towards the bacterial cell surface, where they establish interactions with secondary receptors and ultimately deliver their genetic material into the host cytoplasm. The infection process of flagellotropic phages involves intricate and diverse structural features that enable them to withstand substantial drag forces and torques. In-depth studies have shed light on the molecular mechanisms and dynamic aspects of flagellotropic phages and their interactions with host organisms. Some examples of previously studied phages include the phage F341 [1] that infects *Campylobacter jejuni*, and the phage PBS1 [2] that infects *Bacillus* species. Another example is the phage χ [3], which infects various genera of *Enterobacterales*, presumably by utilizing its tail fiber to bind to the flagellum [4]. Other phages, such as ΦCbK [5], use long capsid fibers to attach to the flagellum and also require active rotation to reach the surface of the host.

*Agrobacterium*, a phytopathogenic bacterium, utilizes horizontal gene transfer to induce crown gall disease in plants [6]. Activation of its virulence gene complex, in response to plant signals, facilitates the transfer of the T-DNA segment from its Ti-plasmid into the plant genome[7]. Among the *Agrobacterium* genus, *Agrobacterium tumefaciens* is extensively studied and capable of infecting a broad range of dicotyledonous plants, including roses, grapes, apples, and tomatoes [8]. Notably, *Agrobacterium* can also infect humans and other animals, particularly those with compromised immune systems, leading to opportunistic infections such as *sepsis*, *monoarticular arthritis*, *bacteraemia*, and *endocarditis* [9]. Moreover, it can produce harmful hydrogen sulfide gas, causing respiratory irritation and damage[10]. *Agrobacterium* has also been linked to certain human diseases, such as cancer and Morgellons disease[11].

Here, we studied the unusual architecture of the lytic flagellotropic phage 7-7-1 and its interaction with its non-infectious host *Agrobacterium* sp. H13-3[12]. Phage 7-7-1 belongs to the *Myoviridae* family[13] and was isolated from compost soil in Germany[14]. This phage contains double-stranded DNA of 247,374 base pairs[15, 16], exhibits high sequence similarity to *Agrobacterium* phage Milano[17] and OLIVR4[18], including an unusual accumulation of cysteine residues in their structural proteins. Typical for *Myoviridae*, it also contains a long contractile tail with multiple elongated and kinked tail fibers[14, 19, 20]. Phage 7-7-1 uses the host lipopolysaccharide as secondary cell surface receptor[21], and its life cycle encompasses an eclipse period of approximately 60 minutes, with complete phage propagation achieved within 80 minutes. The estimated burst size of phage 7-7-1 amounts to 120 particles per bacterial cell[19]. The most unusual characteristic of this phage are the multiple capsid fibers that appear to emerge from the icosahedral capsid. While capsid fibers have been studied in other viruses and phages previously[22–24], they remain highly challenging subjects for structural studies due to their often-flexible nature. The unusual appearance and putative function of the capsid fibers of phage 7-7-1 are highly intriguing, but so far only low-resolution negative stain electron micrograph images are available[19].

To gain insight into the unique structural characteristics of phage 7-7-1, we employed different structural methodologies to investigate its structure, with a focus on the capsid and capsid fibers. Through single particle analysis (SPA) cryo-electron microscopy (cryo-EM), we generated atomic models of the well-organized phage capsid and tail. We determined that the capsid of phage 7-7-1 is composed of a major capsid protein (MCP), a decoration protein (DP) and linking proteins (LP1 & LP2), and that the tail is composed of an outer sheath protein and an inner tube protein. In addition, we determined the presence of capsid fibers, with one individual fiber emerging from each vertex of the capsid. To gain further insight into the structure of these flexible capsid fibers, we employed cryo-electron tomography (cryo-ET). Leveraging machine learning techniques, we successfully used a trained neural network to automatically track the intricate capsid fibers, facilitating quantitative analysis of the tomography data. Our findings highlight the robust attachment capability of the 7-7-1 capsid fibers to flagella during the infection process. Through the synergistic integration of structural, computational, and experimental approaches, we have achieved an in-depth understanding of the structure and functionality of this distinct bacteriophage.

## Results

### Capsid structure of phage 7-7-1

To gain insight into the structural characteristics of the phage 7-7-1 capsid, we choose to use cryo-electron microscopy (cryo-EM) (Data collection parameters in Supplementary Table. 1). Cryo-EM micrographs of phage 7-7-1 reveal the typical structure of this phage: the large icosahedral capsid is coupled to a contractile 135 nm long tail that is intricately adorned with an abundance of bushy tail fibers. A defining characteristic of phage 7-7-1 are the multiple long and curly capsid fibers surrounding each capsid (Supplementary Figure 1, white arrows). Our dataset reveals two distinct variants of the capsid: full capsids containing DNA and empty capsids (Supplementary Figure. 1). Notably, most of the phages observed in the micrographs are intact capsids filled with DNA. To gain detailed insights into the capsid structure, we applied single particle analysis (SPA) and obtained a 3.39 Å structure of the intact, DNA-containing capsids.

The DNA-filled capsid (EMD-52522, Figure 1a) has a diameter of 80 nm when measured from vertex to vertex, and extends 68 nm along the 2-fold symmetry axis. The most striking feature of the 7-7-1 capsid is the presence of capsid fibers, with one fiber emerging from the center of each vertex pentamer (Figure 1b, left). However, due to the flexible nature of these capsid fibers, their densities average out during image processing. Therefore, we could only reconstruct a short section at the fiber base that connects to the capsid, and even these sections could only be resolved at a comparatively lower resolution. The capsid-fiber densities (Figure 1b, green) were visualized in ChimeraX[25] using a small threshold level of 0.01. Furthermore, the capsid belongs to a T = 9 (= 3; = 0; T = + +) triangulation symmetry with a planar outline. (Figure 1c).

**Figure 1.**
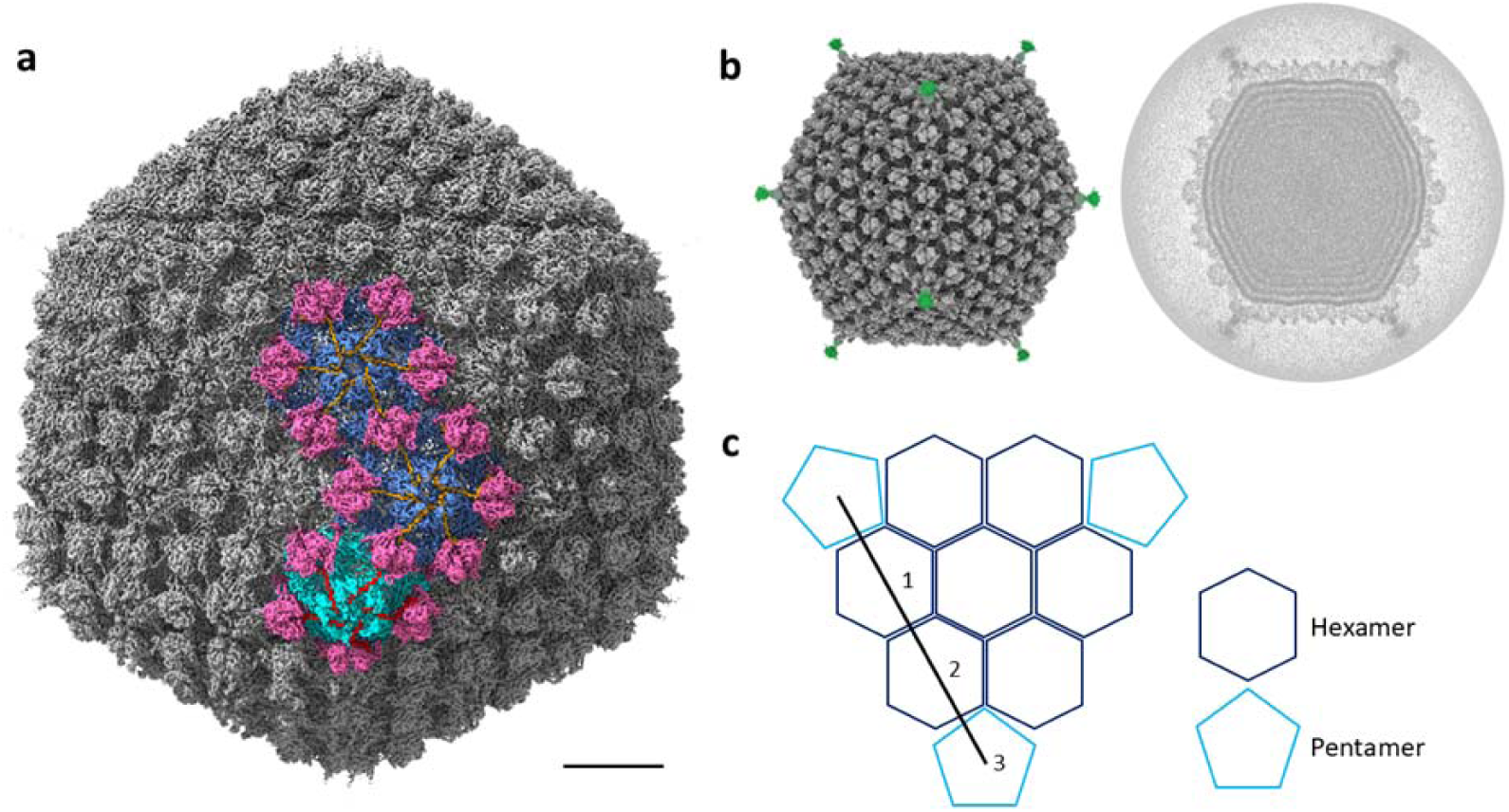
Reconstruction and organization of the DNA containing capsid of phage 7-7-1. **(a)** The icosahedral capsid of phage 7-7-1 filled with dsDNA (EMD-52522) was reconstructed at 3.4 Å resolution. Interlocking hexameric and pentameric capsid structures are highlighted in blue and cyan, respectively. **(b)** Left: the overall cryo-EM density map of phage 7-7-1, revealing the presence of a single capsid fiber at each vertex (green density). This representation used a threshold level of 0.01 in ChimeraX. Right: a plane view of the dsDNA filled inside the capsid. **(c)** Schematic illustrating the 7-7-1 capsid organization, characterized by a T = 9 symmetry. The highlighted line in black signifies the *h* and *k* volumes involved in the calculation of triangulation symmetry, as referenced from the URL (https://viralzone.expasy.org/8577).

In addition to its close relative Milano, the capsid of phage 7-7-1 shows architectural similarity to a range of other phages including *Ralstonia solanacearum* phage GP4[26], satellite phage P2[27], phage N4[28], *Anabaena* phage A-1(L)[29], *Helicobacter pylori* phages KHP30 and KHP40[30], and notably, phage Milano[31], all of which share a common triangulation number of T = 9. Similar to these previously described phages, the icosahedral capsid structure of 7-7-1 is constituted by a lattice of hexamers and pentamers, which are composed of a major capsid protein (MCP) (Figure 1c). This capsid shell contains additional decoration proteins and linker proteins, which play an important role in enhancing capsid stability and potentially contribute to host recognition during the infection process. Specifically, the capsid of 7-7-1 consists of a total of 80 hexamers (Figure 1a, blue) and 11 pentamers (Figure 1a, cyan, with one vertex occupied by the portal complex), thus totaling 535 copies of the major capsid proteins. The MCPs and decoration proteins self-assemble to form the icosahedral shell that acts as the protective envelope for the phage genome.

### Hexamer and pentamer arrangements

The hexamer- and pentamer-forming major capsid protein (gp003, Gene ID: 14012067) of phage 7-7-1 (Figure 2a) consists of 469 amino acids. The residues from A191 to G469 of MCP exhibit sequence and structural similarity to the phage HK97 in an HK97-like fold[32–35]. Structurally, the MCP consists of a C-terminal domain, which constitutes the core, alongside a long loop area and N-terminal loop, which constitutes the peripheral regions of hexamers and pentamers (Figure 2b). The N-terminal residues M1 to A164 of the major capsid protein are not resolved, which may be an indication that they are highly flexible.

**Figure 2.**
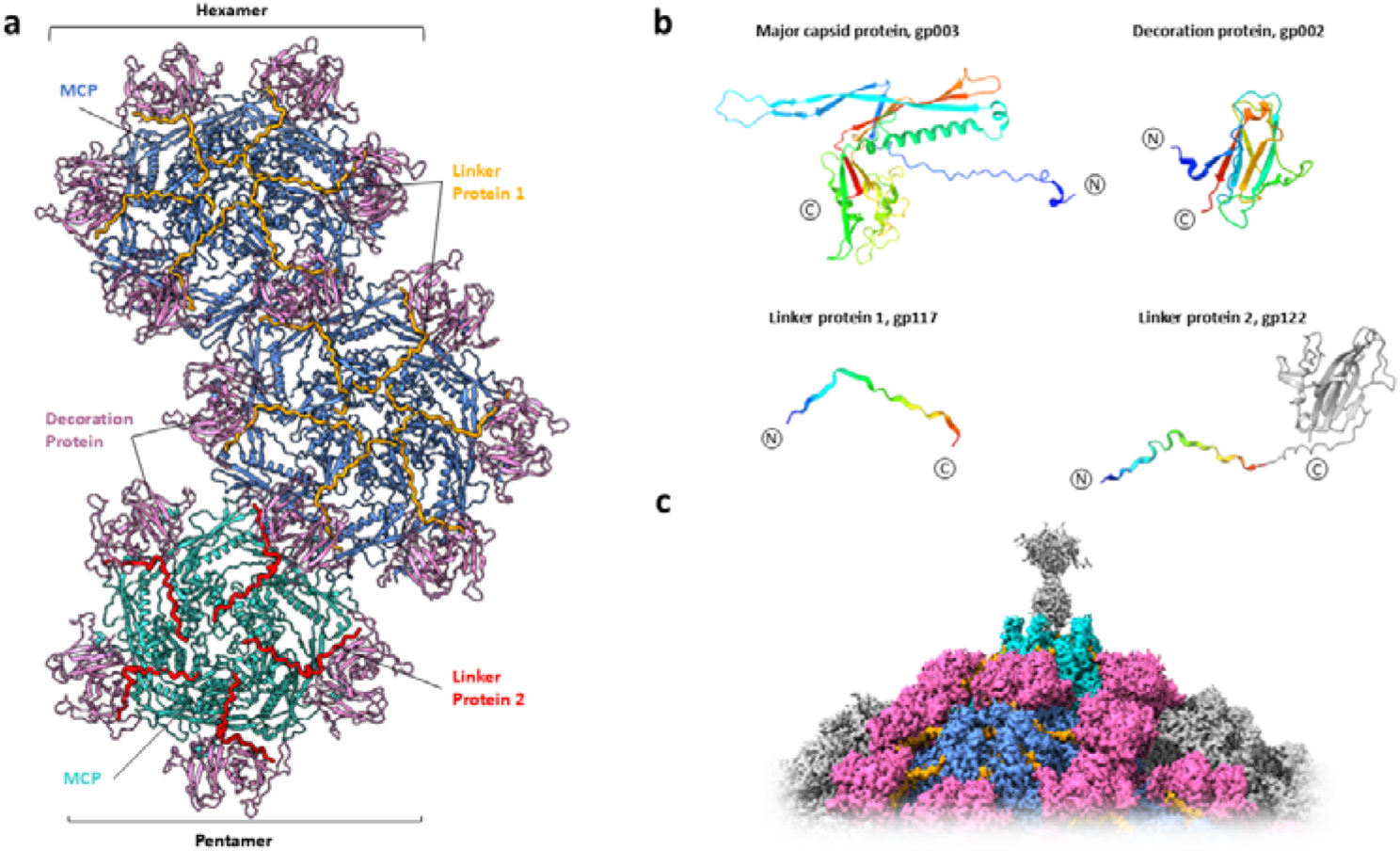
Hexamer and pentamer arrangement in the phage 7-7-1 capsid. **(a)** Structure of pentamer and hexamer differing in their decorating proteins (PDB: 9HZ8). MCP in hexamers are colored blue, MCP in pentamer are colored cyan. The decoration protein, linker protein 1, and linker protein 2 are colored pink, orange, and red, respectively. Each hexamer is decorated by six copies of linker protein 1, while each pentamer is decorated by five copies of linker protein 2. **(b)** Structures of MCP (gp003), decoration protein (gp002), linker protein 1 (gp122), and linker protein 2 (gp117) are shown with N-to C-terminal domains colored in rainbow. **(c)** The capsid-fiber density (gray) protruding from the middle of a pentamer.

The hexametric assembly unit of phage 7-7-1 is composed of six MCP monomers, forming a well-defined structure (Figure 2a, blue). The pentamers are constituted by five MCPs (Figure 2a, cyan). Compared to the hexamers, the pentamers adopt a slightly compressed configuration, accompanied by alterations in their relative orientation, resulting in a more curved arrangement within the pentamer (Supplementary Figure 2). When comparing the hexamers surrounding the pentamer with those not in direct proximity to a pentamer, we observed a subtle augmentation in curvature. These observations provide new structural insights into the organizational dynamics of the capsid assembly of phage 7-7-1.

Hexamers and pentamers are further decorated with a decoration protein (DP, gp002, Gene ID: 14012068) and two linker proteins, namely linker protein 1 (LP1, gp117, Gene ID: 14012070) and linker protein 2 (LP2, gp122, Gene ID: 14012074). These additional proteins exhibit significant sequence and structural similarity to those found in the capsid of phage Milano[31]. The decoration protein comprises 135 amino acids with a β-strand-rich domain (Figure 2b). A dimer of decoration proteins embellishes the capsid surface by bridging the interfaces of pentamers and hexamers. Amongst other interactions, the MCP and the decoration protein share two disulfide bonds: residue C223 of MCP connects to residue C104 of DP, and residue C252 of MCP interacts with C74 of DP.

The assemblies of MCP and DP complexes are further stabilized by linker protein 1 (LP1) and linker protein 2 (LP2), respectively. LP1 (gp117, Figure 2b) on its own forms a relatively unstructured chain of amino acids, but interacts with the capsid facing interface of the DP and extends towards the center of the hexameric MCP complex (Figure 2a). In contrast, LP2 (gp122 Figure 2b) stabilizes the interaction between DPs and pentameric MCP assemblies (Figure 2a). Furthermore, LP2 features a β-sheet Ig-type domain that protrudes the pentameric MCP complex (Figure 2c) into the poorly resolved capsid-fiber density, away from the capsid shell. Ig-type domains are prevalent in phages, and their likely role involves binding to cell surface components, thus tethering the phage to host cells[36–38]. We postulate that five β-sheet Ig-type domains of LP2 form a foundational structure at the center of the pentamer, serving as an initial anchoring point for the capsid fiber. As a whole, these four proteins make up the basic repetitive units that comprise the phage capsid.

### Tail structure of phage 7-7-1

SPA analysis of the tail of phage 7-7-1 revealed its similarity to those of other structurally described *Myoviridae* (Figure 3a, resolution: 3.0 Å). Based on this density map, we observed that the extended tail is 135 nm in length and has a diameter of about 20 nm. The base of the tail is equipped with flexible long tail fibers, components crucial to phage host binding (Supplementary Figure 1a, purple arrows). The phage tail consists of two helically stacked components: the outer sheath (Figure 3b, gp126, Gene ID: 14012078) provides the mechanical force during injection, while a hollow inner tube (Figure 3b, gp127, Gene ID: 14012079) pierces the host membranes during injection and forms a channel for the transfer of genetic cargo[39].

**Figure 3.**
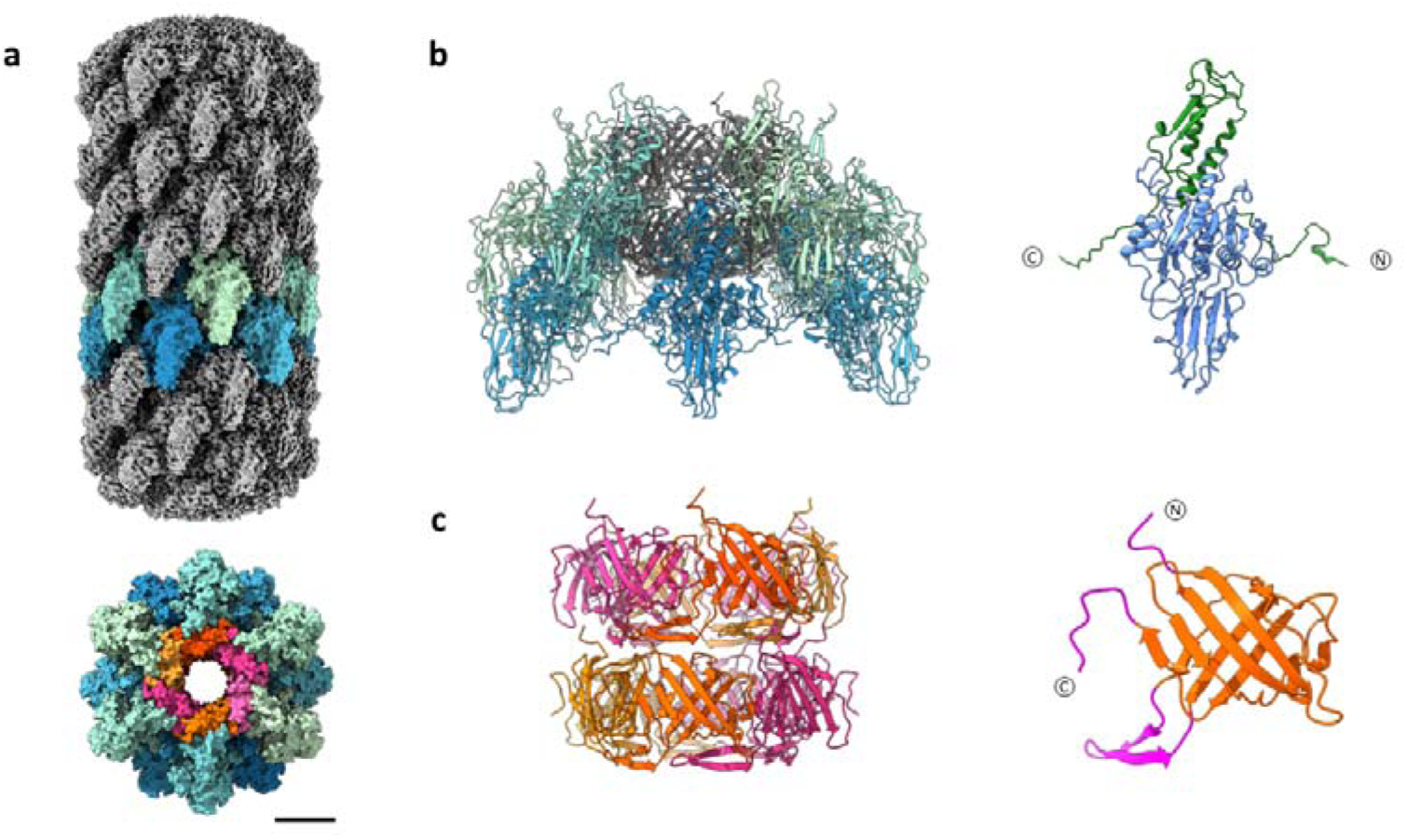
7-7-1 Tail structure arrangement in the phage 7-7-1 tail. **(a)** Top: Side view of 7-7-1 tail density with two sheath subunits rows highlighted (EMD-52521). Bottom: Top view of the 7-7-1 tail structure, the outer sheath and inner tube colored with a gradient of blue-green and orange-pink, respectively (scalebar: 6nm). **(b)** Left: Ribbon diagram of the tail complex model (PDB: 9HZ7). Sheath subunits are highlighted. Right: The ribbon model of 7-7-1 sheath protein gp126. Interaction domains highlighted in green (residues 1-25 and 385-500) **(c)** Left: Organization of the tail tube complex. Right: The ribbon model of 7-7-1 tube protein gp127. The interprotein interaction domains, N-terminus (residues 1-7), C-terminus (residues 128-135), and loop (residues 43-61) are highlighted in magenta.

The stacked sheath proteins form a C6-symmetric helical interlocking structure. Every 7-7-1 sheath subunit has two long arm extensions at the N- and C-termini. Accompanied by two neighboring subunits, each arm contributes to the formation of an interlocking β-sheet motif (Figure 3b). This in turn leads to a helical assembly of the tail sheath complex (twist of 27.35° and a rise of 35.25 Å). Furthermore, the tail sheath complex has a lumen of approximately 70 Å in which the inner tube resides. The 7-7-1 inner tube structure also exhibits a C6 symmetry, aligning with the surrounding sheath proteins (Figure 3c). Tube protein subunits oligomerize into hexameric ring-like structures that stack on top of one another, creating a hollow β-barrel tube structure with an inner diameter of approximately 40 Å. The three main interaction domains that stabilize the tube assembly (the N-terminal, C-terminal, and loop interaction domain) are highlighted in pink. The combined sheath and tube complex is held together by a multitude of hydrogen bonds and electrostatic interactions. The inner surface of the sheath assembly is positively charged, while the outer surface of the tail tube is negatively charged (Supplementary Figure 3). Taken together, the structural investigation of the 7-7-1 phage tail provided us with insights into the intricate organization of the sheath- and tube proteins, making up an important part of the infectious machinery of phage 7-7-1.

### Cryo-electron tomography and segmentation of phage 7-7-1

Phage 7-7-1 was further investigated by cryo-ET to investigate the structurally flexible components of phage 7-7-1, as well as to gain insight into host-phage interactions during the initial infection process. In a representative cryo-ET image (Figure 4a) of the bacteria-phage 7-7-1 mixture, we found an accumulation of phage particles in close proximity to a cluster of flagella. Notably, the capsids of phage 7-7-1 were oriented toward the flagella, indicating their capability to establish connections with the flagella via structures located on their capsids. The accumulation of full-DNA phages around the flagella provided evidence of initial phage-host interaction of 7-7-1. Furthermore, analysis of individual phages in the tomograms revealed that the capsids, while in close proximity, appear not to be in direct contact with the flagella. This suggests that the interaction of the phage with the bacterial flagellum involves the filaments located on the capsid.

**Figure 4.**
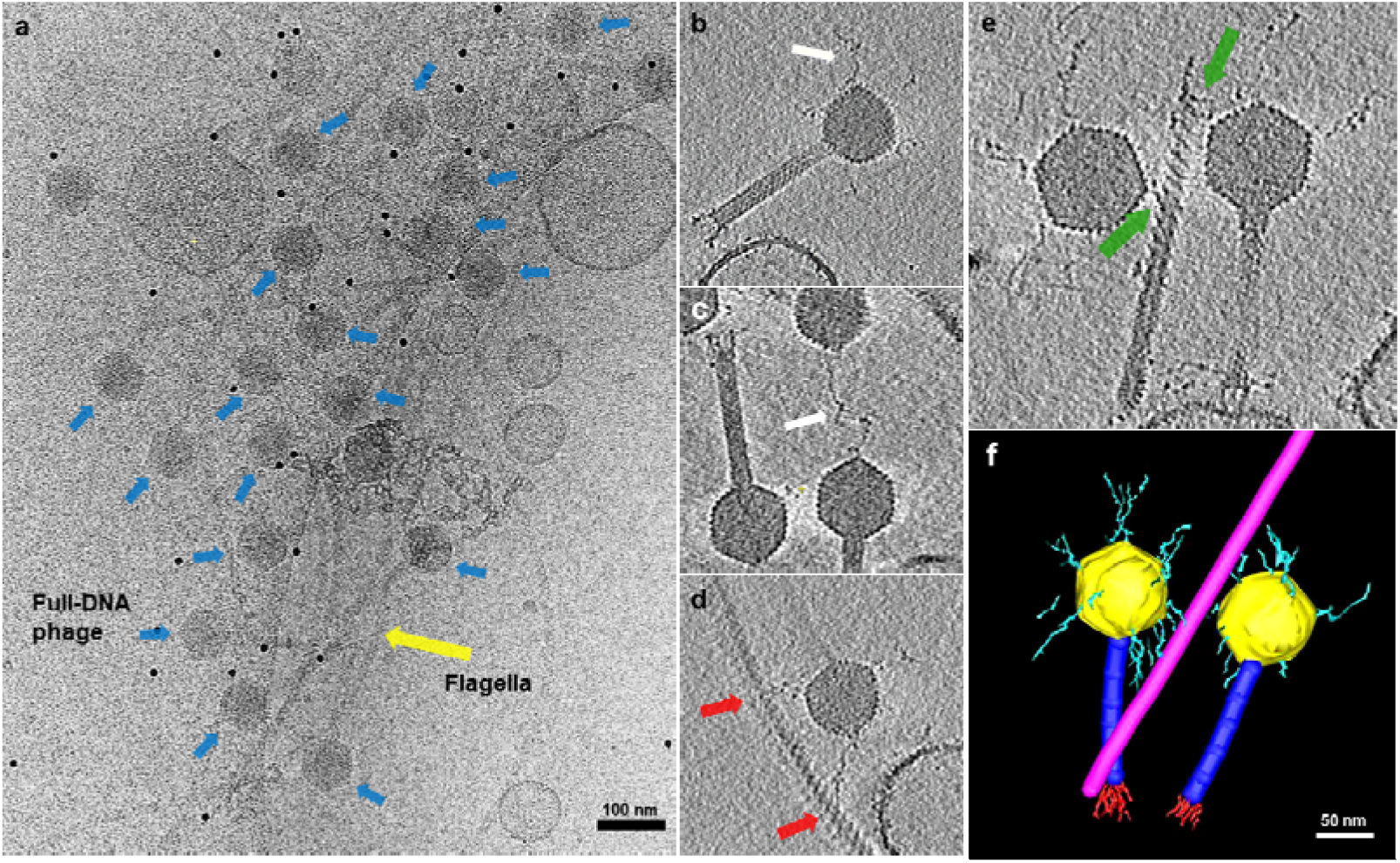
Cryo-electron tomography and segmentation of phage 7-7-1 attached to *Agrobacterium* flagellum. **(a)** A representative cryo-ET cross-section (slice) is presented, revealing multiple instances of full-DNA phage 7-7-1 (blue arrows) attached to a cluster of flagella (yellow arrow). Scale bar: 100 nm. The tomographic data clearly reveal the presence of the characteristic icosahedral capsid, contractile tail, tail fibers, and capsid fibers. **(b)** Example of long flexible fibers (white arrow) of a full-capsid phage 7-7-1, originating from the capsid vertices. **(c)** Occasionally, it can be observed that the flexible capsid fibers are connecting two phages (white arrow). (**d**) The interaction of two phage capsid fibers with the host flagellum (red arrows). (**e**) Two phages attached to the same flagellum (green arrows), further emphasizing the strong interaction between phage 7-7-1 and the host flagellum. **(f)** A volumetric representation of the tomogram subregion shown in **e**, with manual segmentation performed using IMOD. Scale bar: 50 nm.

To investigate the architecture and function of the flexible capsid fibers during the various stages of the infection process, we focused on complete phage 7-7-1 particles interacting with its host. More specifically, we explored phage 7-7-1 in conjunction with its host flagellum, leading to the identification of four distinct states: free particles with and without DNA and adsorbed particles with and without DNA. These selected particles represent the different stages during phage infection, including free phages (state 1), phages during the early-infection stage once attached to the host (state 2), post-infection (empty) phages still attached to the cell (state 3), and post-infection (empty) free phages (state 4).

The pre-infection phages (state 2) filled with DNA were used to study the interaction between capsid fibers and flagella. Sub-volumes of individual full-capsid phages were extracted from the tomograms. They confirmed the presence of long, flexible fibers connecting the capsids with the flagellar filament (Figure 4b, 4c, 4d, 4e). More specifically, one long, flexible fiber emerges from each of the vertices of the 7-7-1 capsid. Our SPA result has revealed that the fibers emerge from the 11 pentamers on the capsid. However, due to the missing wedge artifact in the tomograms, which is caused by the limited tilt range of the specimen holder and blocks full sample illumination, and the inherent flexibility of the fibers, not all capsid fibers were fully visible in the tomograms. Furthermore, accurate measurements of the fiber length posed challenges due to their flexible and curly nature.

Intriguingly, we found that the capsid fibers of phage 7-7-1 not only seem to interact with the host’s flagellum but could also interact with fibers from other 7-7-1 capsids (Figure 4c, white arrow). However, the nature of the interaction between the fibers and their potential role to aid each other’s probability of success for infection remains unknown. At present, the proteins, their functions, and the interaction mechanisms of the head fibers with host flagella are also unknown. Previous studies have identified three candidate receptor binding proteins (RBPs), gp4, gp102, and gp44, based on their ability to bind to host cells[15]. It remains to be seen whether these three putative RPBs specifically interact with host flagella. Figure 4d further highlights the intriguing observation of one flagellum attached to two 7-7-1 phages by capsid fibers, suggesting the possibility of multiple phages concurrently attaching to the same flagellum via multiple capsid fibers. A closer inspection of the helical flagellar architecture depicted in Figure 4c, 4d, and 4e reveals that the phage capsid fibers wrap around the flagellum. This observation corresponds to the established model describing the interaction between the capsid fiber of bacteriophage [CbK and flagella of its host[40].

Due to the intricate complexity and heterogeneous nature of the capsid fibers, manual segmentation using the IMOD segmentation function[41] was chosen for initial analysis. Consequently, the sub-tomogram presented in Figure 4d was manually segmented in IMOD and revealed multiple capsid fibers attached to the flagellum (Figure 4f). These findings shed light on the complex interactions between phage 7-7-1 and its host’s flagella, underscoring the importance of such connections in the infection process.

### Capsid fiber analysis using machine learning

To streamline the labor-intensive process of manually segmenting capsid fibers, we applied our previously developed neural network to automate the detection of tail fibers in reconstructed tomograms of phage [Kp24[42]. Here, this network was specifically trained using manual segmentations of capsids, capsid fibers, and flagella, and subsequently optimized through multiple iterations. Once trained, the network was applied to all tomograms, resulting in the generation of comprehensive 3D maps that accurately captured the structures of capsid fibers and flagella. This approach significantly reduced the time and effort required for fiber detection and facilitated efficient analysis of the phage structures. These results illustrate the structural attachment of phages via their capsid filaments to the flagellum during the initial infection stage.

As mentioned above, we defined the pre-infection phages based on four key characteristics: (1) the presence of a full capsid, (2) attachment of capsid fibers to flagella, (3) intactness of the phage structure, and (4) absence of contact with a bacterial cell. After extraction and format conversion, 13 capsid fiber structures of pre-infection phages were generated. Through visual inspection, we carefully selected and presented 6 groups of representative trimmed tomograms and their corresponding capsid fiber structures in Supplementary Figure 4. These 3D structures exhibit distinct conformations for each capsid fiber, highlighting the dynamic nature of these components.

Following an extraction and format conversion process, we successfully generated 13 pre-infection phage 7-7-1 structures that were attached to the flagellum. Measuring the length of capsid fibers within these structures revealed a range between ∼ 88 nm to 114 nm. While the flexibility and curvature of capsid fibers posed challenges to obtain precise measurements, these findings demonstrate the remarkable capability of phage 7-7-1 to establish a robust attachment but at a considerable distance between the capsid and the flagellum.

In figure 5, we present two representative 3D structures of capsid fibers from phage 7-7-1 attached to the flagellum. These 3D structures exhibit distinct conformations for each capsid fiber, highlighting their dynamic nature=. Particularly, the depicted structures in Figure 5b (capsid fibers indicated with white lines) illustrate the capsid fibers extending from the phage and enveloping the flagellum. This mode of fiber-flagellum interaction closely resembles the well-established interaction observed between the capsid fiber of bacteriophage [CbK and its host flagellum[40].

**Figure 5.**
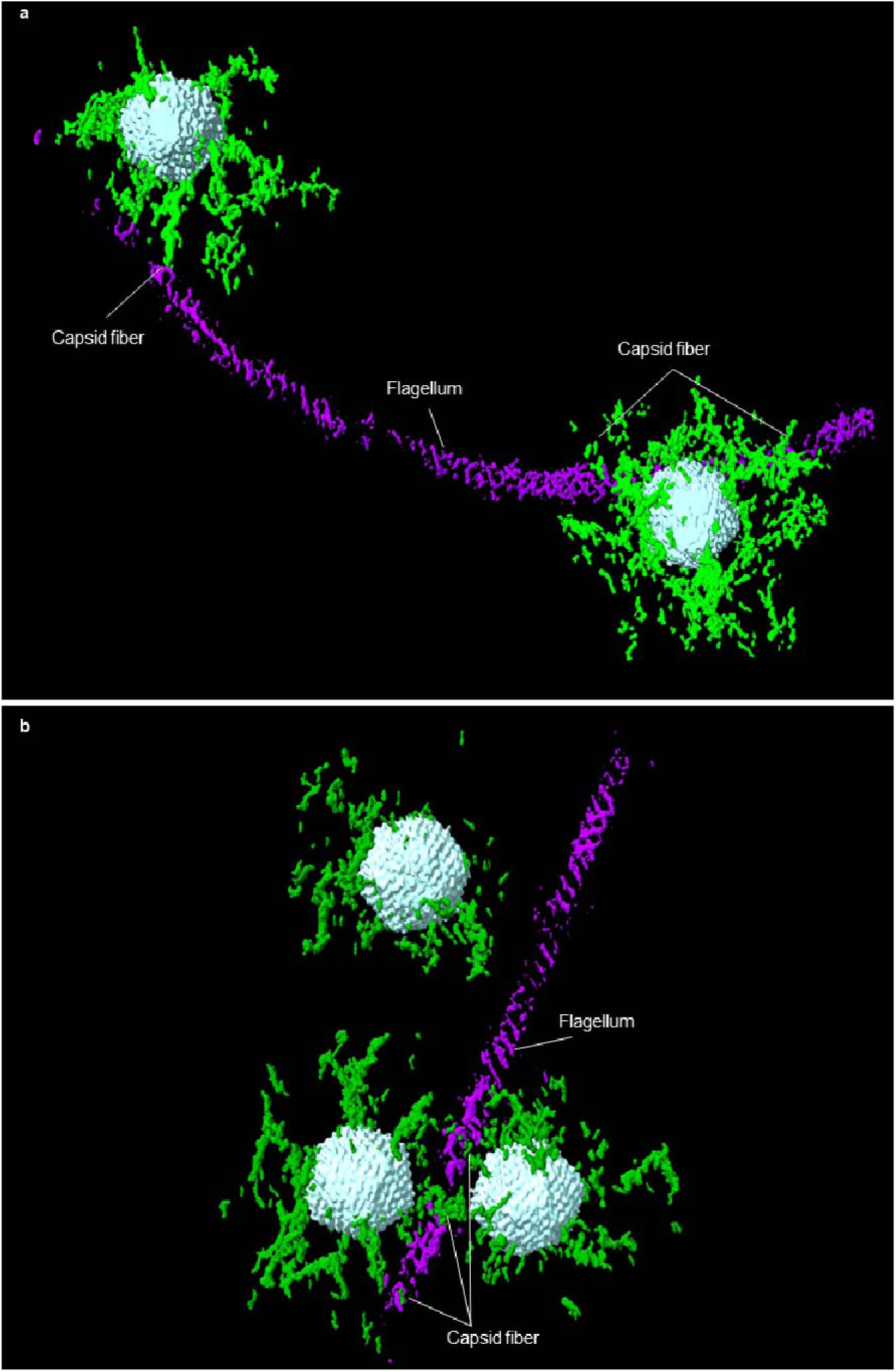
3D structures of capsid fibers of phage 7-7-1 attached with flagellum. The 3D structures of capsid (cyan), capsid fibers (green), and flagellum (purple) were generated using the neural network. Flagellum-associated phage 7-7-1 particles are oriented with their capsids toward and close to the flagellum. Structures of phage 7-7-1 with capsid fibers extending from the phage and wrapping around the flagellum (white lines mark capsid fibers).

### Effect of Proteinase K degradation on capsid fiber

Proteinase K treatment was utilized to determine if the capsid fibers of bacteriophage 7-7-1 are formed by protein. Proteinase K was added at a final concentration of 1% to the concentrated phage 7-7-1 stock and incubated for 20, 40, and 60 minutes at 37 °C. Following cryo-EM imaging, the micrographs revealed a progressive reduction in the number of capsid fibers with increased incubation time (Figure 6a), indicating the time-dependent degradation of the capsid fibers upon Proteinase K treatment. Spot assays of Proteinase K-treated phage samples demonstrated that visible host cell lysis was achieved over time. While lysis occurred at a 10^-6^ dilution of the untreated phage stock, a 10^1^ and 10^4^ higher titer was required to obtain lysis with the Proteinase K-treated phage samples after 40 and 60 min, respectively (Figure 6b). This result coincides with the time-dependent degradation of capsid fibers. In conclusion, Proteinase K effectively degrades capsid fibers and negatively affects infectivity of phage 7-7-1.

**Figure 6.**
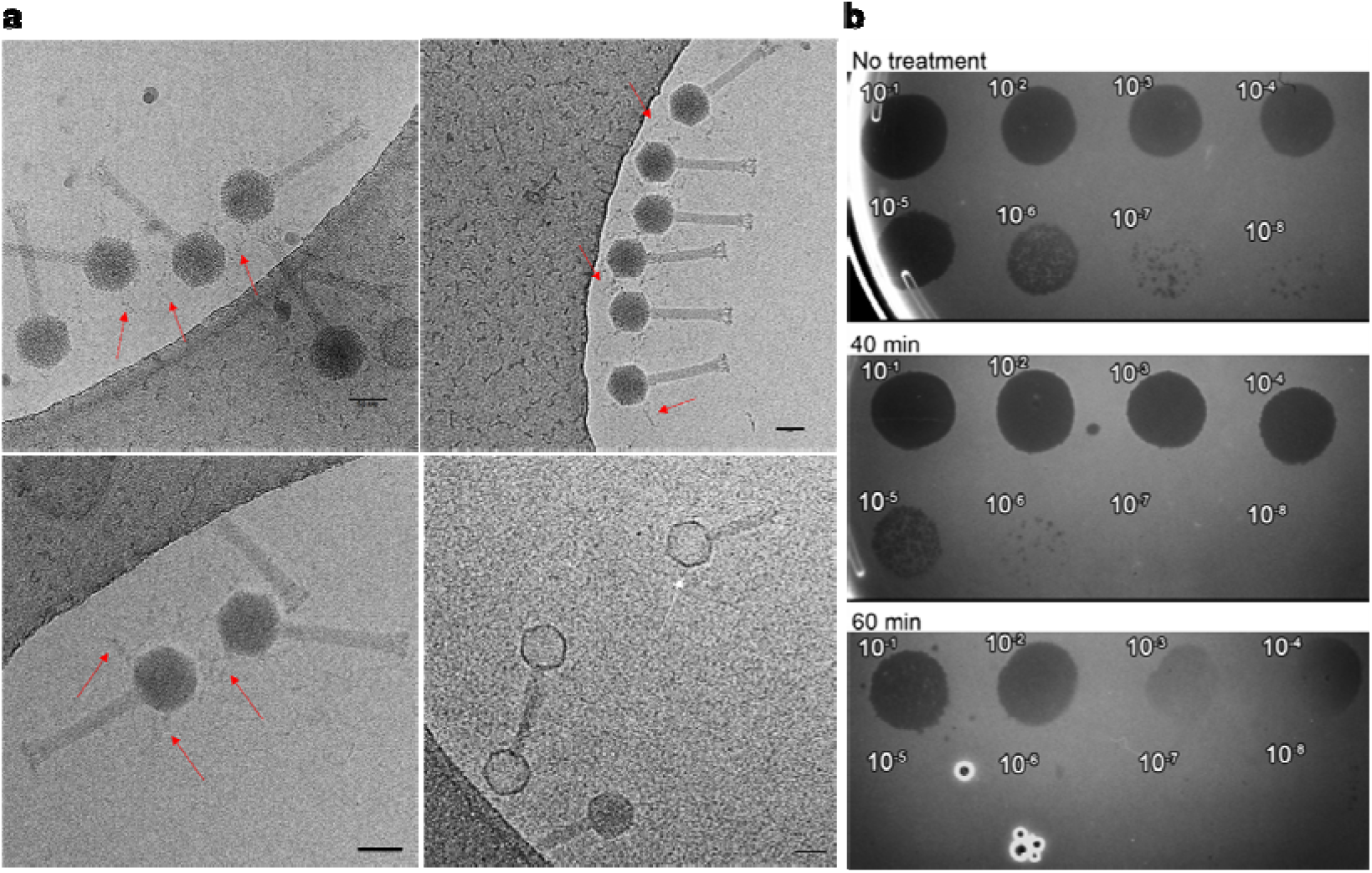
Effect of Proteinase K degradation on capsid fiber and infectivity over time. **(a)** Representative cryo-EM micrographs showing the degradation of the capsid fibers of bacteriophage 7-7-1 after no treatment (top left), 20 minutes (top right), 40 minutes (bottom left), and 60 minutes (bottom right) of Proteinase K treatment. Scale bar depicts 50 nm. Red arrows highlight the capsid fibers, the white arrow highlights a small part of remaining capsid fiber after 60 minutes of treatment. **(b)** Spot assays of serially diluted phage 7-7-1 on a lawn of its host strain without treatment, and after 40 minutes and 60 minutes of Proteinase K treatment.

## Discussion

Capsid fibers assume a critical role in facilitating the attachment of viral capsids to host receptors. Various phages and viruses exhibit intriguing variations in the structural characteristics of these capsid fibers. For instance, *Bacillus subtilis* phage φ29[43] contains capsid fibers that are attached to quasi-3-fold symmetry positions on the capsid, characterized by a protruding stem without a terminal sphere[23, 44]. Other phages, such as *Caulobacter crescentus* phage φCbK and *Vibrio alginolyticus* phages φSt2 and φGrnl feature only a solitary flexible fiber positioned atop the viral head[40, 45, 46]. Some *Vibrio* phages possess fibers, often referred to as antennae, terminated by trilobate structures anchored on the viral head[47]. Another form of capsid fibers can be found in the *Agrobacterium tumefaciens* phage Atu_ph07, which resemble thin hair-like fibers[48]. Here we determined a novel arrangement of capsid fibers in phage 7-7-1, with one fiber emerging from each vertex. More specifically, the fibers emerge from each pentamer of the capsid, totaling in 11 capsid fibers. The final (12^th^) vertex is occupied by the phage-tail connection.

Despite the new structural insights into the unusual capsid and capsid fibers of phage 7-7-1 reported here, many questions remain. For example, the structure of the 7-7-1 capsid fiber is still unknown, and we can only suggest that the five β-sheet Ig-type domains of link protein 1 gp122 form a foundational structure at the center of the pentamer, serving as an initial anchoring point for the capsid. Drawing an analogy to the fiber proteins found in human adenoviruses, which are also positioned at each vertex of the capsid, we postulate that 7-7-1 capsid fibers might share a common architectural framework. According to this model[49], the capsid fiber protein can be envisioned as a trimer, with each monomer comprising an N-terminal tail, a central shaft consisting of repetitive sequences, and a C-terminal globular knob domain. In this context, the N-terminal ∼45 residues of the fiber protein are notably conserved across various phages and are primarily responsible for binding to the penton base[50, 51]. On a structural level, within one capsid fiber, a fiber protein trimer potentially binds to a pentameric complex located at the fivefold symmetry axes of the capsid. For an individual monomer within the trimeric protein, approximately 11 residues of the N-terminal tail form stable contacts by inserting between two adjacent penton monomers[52], which means there is a stable attachment between the fiber and the virion when other regions of the fiber move[53, 54]. By featuring 11 capsid fibers, phage 7-7-1 circumvents steric limitations presented by the relatively flat capsid surface. This architectural adaptation likely enhances the phage’s ability to bind to flagella in random orientations and at different rotational speeds, which may improve the likelihood and efficiency of attachment.

Deciphering the unique morphology of the 7-7-1 capsid presents a notable challenge for structural investigations. The considerable size of the capsid and the heterogeneity of the capsid fibers introduce complexities in both data collection and analysis. They require the integration of diverse complementary structural techniques to achieve a comprehensive understanding of the overall structure.

The building components of the capsid in phage 7-7-1 are the MCP gp003, decoration protein gp002, and two linker proteins linker protein 1 gp117 and linker protein 2 gp122. Typically, phage capsids are composed of one or two MCPs along with decoration proteins that assist in linkage and reinforce the MCPs interactions[32]. Similarly, the capsid of 7-7-1 is also assembled by MCPs and additional proteins. This systematic arrangement ensures a stable and secure encapsulation of the genome inside the capsid, contributing to the phage’s functional efficacy. The tail proteins of phage 7-7-1 are highly conserved amongst *Myoviridae* [55]. For instance, the 7-7-1 sheath protein gp126 shares 90% of sequence identity with the Milano sheath protein gp20 and has a closely related morphology[56], thus sharing a similar contractile tail-like injection system.

While previous work has only indicated the presence of capsid head fibers in low resolution EM images, we gained more detailed insight into the structural importance of head fibers in the infection process of phage 7-7-1. Due to the challenging flexible nature of these fibers, little structural and functional insight was available. Here we used cryo-ET to image the host and phage 7-7-1 in 3D and at macromolecular resolution. Initial imaging revealed that the capsids of phage 7-7-1 were oriented toward flagella but at a considerable distance. This already suggested their ability to establish direct connections with the flagella via fibers situated on their capsid surfaces. Furthermore, our data showed that multiple phages can attach to a single flagellum, which could indicate a collaborative association potentially facilitated by the flagella themselves.

Surprisingly, we found that phage 7-7-1 can also form connections with other phages via their head fibers. We propose that phage 7-7-1 could act collaboratively to accomplish successful infection: an initial phage seeks out and attaches to a flagellum, and subsequently, another phage can locate and connect to the first phage via the capsid fibers, thus facilitating its proximity to the bacterial cell and subsequent completion of infection. The interaction details between head fibers are still unknown and need further investigation in future. In addition, we observe the adsorption of multiple capsid fibers to the flagellum, illustrating the robust attachment capability of the 7-7-1 capsid to flagella.

Drawing an analogy to the capsid fibers observed in phages [Cb13 and [CbK[40], which are also positioned at each vertex of the capsid, we hypothesize that the capsid fibers of 7-7-1 may share a common structure and employ a similar interaction mechanism with the flagella of the host cell. It is plausible that phage 7-7-1 initially adheres to the flagellum through the head filament, subsequently leading to an irreversible attachment of the phage tail to the lipopolysaccharide (LPS) layer of the host^21^, which serves as secondary cell surface receptor. Consequently, we surmise that the interaction between the head filament of phage 7-7-1 and the flagellum resembles that observed between the [CbK phage’s head fiber and the flagellum[45, 57]. In this model, a portion of the phage’s head fiber encircles the flagellum, facilitating the localization of bacteriophages to the cell surface as the flagellar filament rotates. Subsequently, the tail of phage 7-7-1 is anticipated to bind to the LPS culminating in irreversible attachment and DNA injection[21].

## Materials and methods

### Sample preparation

Protocol for propagating and purifying phage 7-7-1 was followed as previously described[21, 58]. Phage 7-7-1 was concentrated as followed: 1000 µl of phage 7-7-1 preparation (1×10_12_ pfu/mL) was added to a Centricon with a 10-kDa cellulose filter and centrifuged 2-3 times at 5,000 × g for 15 min each to approximately xx μl until no buffer eluted from the protein concentrator.

For SPA reconstruction, the samples were plunge frozen using the Leica EM GP: 3.0 µl of the concentrated phage was added to a glow discharged Quantifoil R2/2, 200 mesh Cu grid, incubated for 10 s at 20 °C with approximately 85% relative humidity, pre-blotting time was 30 s, blotted for 0.7 s, and automatically plunged into liquid ethane.

### Proteinase K treatment

Two microliters of Proteinase K (10% w/v) were added to 18 µl of the concentrated phage 7-7-1 suspension and incubated at 37 °C for 20, 40, and 60 minutes. For subsequent imaging, 3 µl of the mixture was added to a glow discharged Quantifoil R2/2, 200 mesh Cu grid, incubated for 10 s at 20 °C with approximately 85% relative humidity, pre-blotting time was 30 s, blotted for 0.7 s, and automatically plunged into liquid ethane. All the vitrified samples were stored in liquid nitrogen until use.

To determine the infectivity of Proteinase K treated phage, 90 μl of 7-7-1 stock at 5.5 x 10^10^ pfu/ml were mixed with 10 μl of 10% (w/v) Proteinase K in H_2_O. This mixture was incubated in a 56 °C heat block for 60 min alongside a control sample that did not receive Proteinase K, and samples were withdrawn at 40 and 60 min. Next, phenylmethylsulfonyl fluoride (PMSF) was added to a final concentration of 5 mM to stop proteolysis. After incubation at 4 °C for 24 hours, each sample was diluted to a final volume of 500 μl using TM buffer. Samples were transferred to Microcon centrifugal concentrators (100,000 MWCO) and centrifuged at 14,000 x g until 90% of liquid had flowed through the membrane. Membranes were washed five times by adding 450 μl of TM buffer followed by centrifugation. The Microcon column was inverted in a fresh collection tube and centrifuged for 1 min at 1,000 x g to collect the phage sample.

### Phage spot assays

Spot assays were used to estimate the titer of infectious phage particles in Proteinase K treated samples. TYC medium containing 0.5% agar (TYC soft agar) was melted and cooled to 50 °C. *Agrobacterium* sp. H13-3 was grown in TYC until an OD_600_ of 0.3 was reached. Motility was verified by phase contrast microscopy. Next, 100 μl of bacterial culture were mixed with 4 ml molten TYC soft agar and poured onto the surface of a TYC agar plate. After the soft agar had solidified, 10 μl of serial 7-7-1 dilutions in TM buffer were spotted on the surface of the plate. The formation of clearing zones after 24 hours of incubation at 30 °C was indicative of phage-mediated lysis.

### Imaging conditions for SPA

Phage 7-7-1-containing grids were clipped and loaded into a Titan Krios electron microscope (Thermo Fisher Scientific, TFS) operated at 300 kV, equipped with a K3 direct electron detector and BioQuantum energy filter using a slit width of 20 eV (Gatan, Inc). Movies were recorded using SerialEM[59] in super-resolution mode at 64,000 nominal magnification, corresponding to a calibrated pixel size of 0.685 Å, with a defocus range of −1 to −2.5 µm, and a total dose of 40 e/Å _2_(see data collection parameters in Supplementary Table. 1).

### SPA data processing

Data processing was performed using CryoSPARC[60]. Micrographs were imported, patch motion corrected and patch CTF estimated. For the reconstruction of the phage capsid, particles were manually picked for template generation and subsequent template picking. After multiple rounds of 2D classifications, the best classes were submitted to ab-initio model generation using icosahedral symmetry. The resulting best model was used for homogeneous reconstruction, with local and global CTF corrections, as well as Ewald sphere correction. The particles were then submitted to reference-based motion correction and a final round of homogenous reconstruction as described before, yielding the final phage capsid reconstruction. For the reconstruction of the phage tail, particles were also manually picked and used for template generation and subsequent filament tracing (filament diameter 220 Å, separation distance 0.5 diameters, minimum length 4 diameters). The resulting picks were used in multiple rounds of 2D classification and the best classes were used for ab-initio model generation. The best model was analyzed with the symmetry search utility and yielded an estimated 27° twist and a 35 Å rise. Next, helical refinement was performed using these parameters together with C6 symmetry, yielding the final phage tail reconstruction. Map-map fourier shell correlation curves of both maps (cut-off 0.143) are shown in Supplementary Figure 5.

### Model building

ModelAngelo[61] was used for initial model generation without providing sequence input. Subsequently, protein sequences were identified with findMySequence[62] referenced against the partially annotated phage 7-7-1 genome[17] and fed back into ModelAngelo. The sequence-based model was then analyzed in ChimeraX[25] and any missing regions were added with the Modeller package[63]. The models were then interactively improved using ISOLDE[64] and refined in Phenix[65]. The model building statistics are available in Supplementary Table 2.

### Sample preparation for cryo-ET

*Agrobacterium* sp. H13-3 strain was grown overnight in TYC (0.5% tryptone, 0.3% yeast extract, and 0.087% CaCl_2_ x 2H_2_O [pH 7.0]) at 30 °C in a shaking incubator at 180 rpm/min. Two hundred microliters of bacterial cell cultures (0D_600_ = 0.2) were sedimented at 3,000 × g for 15 min at room temperature and suspended in 15 µl motility buffer (0.5 mM CaCl_2_, 0.1 mM EDTA, and 20 mM HEPES, pH 7.4). Subsequently, the concentrated cell suspension was mixed with 15 µl of phage 7-7-1 (1×10_12_ pfu/mL) and 2 µl 10-nm gold beads (Cell Microscopy Core, Utrecht University, Utrecht, The Netherlands). The mixture was incubated at room temperature without shaking for 3-10 min prior to plunge freezing. Using the Leica EM GP (Leica Microsystems, Wetzlar, Germany), 3.8 µl of the bacteria, phage and gold bead solution (volume ratio: 15:15:2) was applied to a glow discharged Quantifoil R2/2, 200 mesh Cu grid (Quantifoil Micro Tools GmbH, Jena, Germany), which was incubated for 30 s prior to blotting at 20°C with approximately 95% relative humidity. The grids were blotted for 1 s and automatically plunged into liquid ethane. Vitrified samples were transferred to storage boxes and stored in liquid nitrogen until use.

### Imaging conditions of cryo-ET

The grids containing *Agrobacterium* sp. H13-3 bacteria and phage 7-7-1 were clipped and loaded into a Titan Krios (Thermo Fisher Scientific (TFS)) transmission electron microscope equipped with a K3 BioQuantum (Gatan, Inc) direct electron detector operating in counting mode and the energy filter set to a 20 eV slit. Targets were chosen based on the presence of flagella attached to bacterial cells that were located in a hole of the carbon film of the EM grid. Cryo-EM micrographs of phage 7-7-1 attached to flagella were collected using SerialEM set to a dose symmetric tilt scheme between -54° and 54°, with 2° tilt increments[59, 66]. The selected nominal magnification was 26,000, which corresponds to a pixel size of 3.28 Å. The defocus of the collected data ranged from -4 to -6 µm. The total dose per tilt series was 100 e-/Å.

### Cryo-ET data processing

Motion correction, tilt series alignment using gold fiducials, and tomogram reconstruction was carried out using IMOD[41]. The datasets were binned by a factor of 2. From the reconstructed tomograms, regions showing phages interacting with flagella were selected for segmentation using IMOD. Visualization of data was performed in IMOD and Fiji[67].

### Fiber analysis using machine learning

A 100-layer mixed-scale dense neural network was trained to detect the capsid fibers of phage 7-7-1 in reconstructed tomograms. The training setup was similar to that used in other studies utilizing mixed-scale dense networks[68–70]. For the training process, manual segmentations of the tail fibers from 9 phages were utilized. The ADAM algorithm[71] was employed to minimize the cross-entropy loss, with random rotations and flips applied for data augmentation. The entire training process took a few hours and was halted when no significant improvement in the loss was observed. After training, the network was applied to all tomograms, yielding 3D maps of the fiber structures. To perform additional analyses, manual annotation was conducted on the tomograms for each phage, identifying the center of the fiber structure and the position of the head. Using this annotated information, a 3D map of the fiber structure for each phage was extracted from the 3D fiber maps, resulting in individual maps for 13 capsid fiber structures of pre-infection phage. To ensure proper alignment, the maps were aligned to each other using both the manually annotated position information and subsequent automatic alignment through maximizing cross-correlation. The extracted structures were visualized using ChimeraX for further examination and analysis.

## Acknowledgements

R.O. was supported by the China Scholarship Council (CSC) with project number 201906280465. This work benefited from access to the Netherlands Centre for Electron Nanoscopy (NeCEN) at Leiden University, which was funded in part by the Netherlands Electron Microscopy Infrastructure (NEMI), project number 184.034.014 of the National Roadmap for Large-Scale Research Infrastructure of the Dutch Research Council (NWO). This work was funded by the National Science Foundation fund number IOS-2054392 to BES. We thank Floricel Gonzalez and Abigail Horton for 7-7-1 phage preparations.

## Contributions

Conceptualization: A.B.; methodology: A.B., A.S., N.C.E., R.O., T.H., W.N..; formal analysis: A.S., N.C.E., R.O., T.H., W.N. investigation: A.S., N.C.E., R.O., T.H., W.N.; resources: A.B., B.E.S., D.M.P.; data curation: A.B., A.S., B.E.S., N.C.E., R.O., T.H., W.N.; writing original draft: A.B., A.S., R.O., T.H., W.N.; writing— review and editing: all authors; supervision: A.B.; project administration: A.B.; funding acquisition: A.B., A.S., B.ES., D.M.P., R.O.

## Data availability

The 3D cryo-EM maps generated in this study have been deposited at EMDB (the Electron Microscopy Data Bank, https://www.ebi.ac.uk/emdb/) with accession code EMD-52522 for the full capsid for phage 7-7-1, and the code EMD-52521 for the tail of phage 7-7-1. Protein prediction data reported in this paper will be shared by the lead contact upon request. The atomic coordinates generated in this study have been deposited at wwPDB (Worldwide Protein Data Bank, http://www.wwpdb.org/) with accession PDB ID: 9HZ8 for pentamer-hexamers of the full capsid of phage 7-7-1, and the accession PDB ID: 9HZ7 for the tail (2 layers) of phage 7-7-1.

## Supplementary Materials

**Supplementary Table 1.**
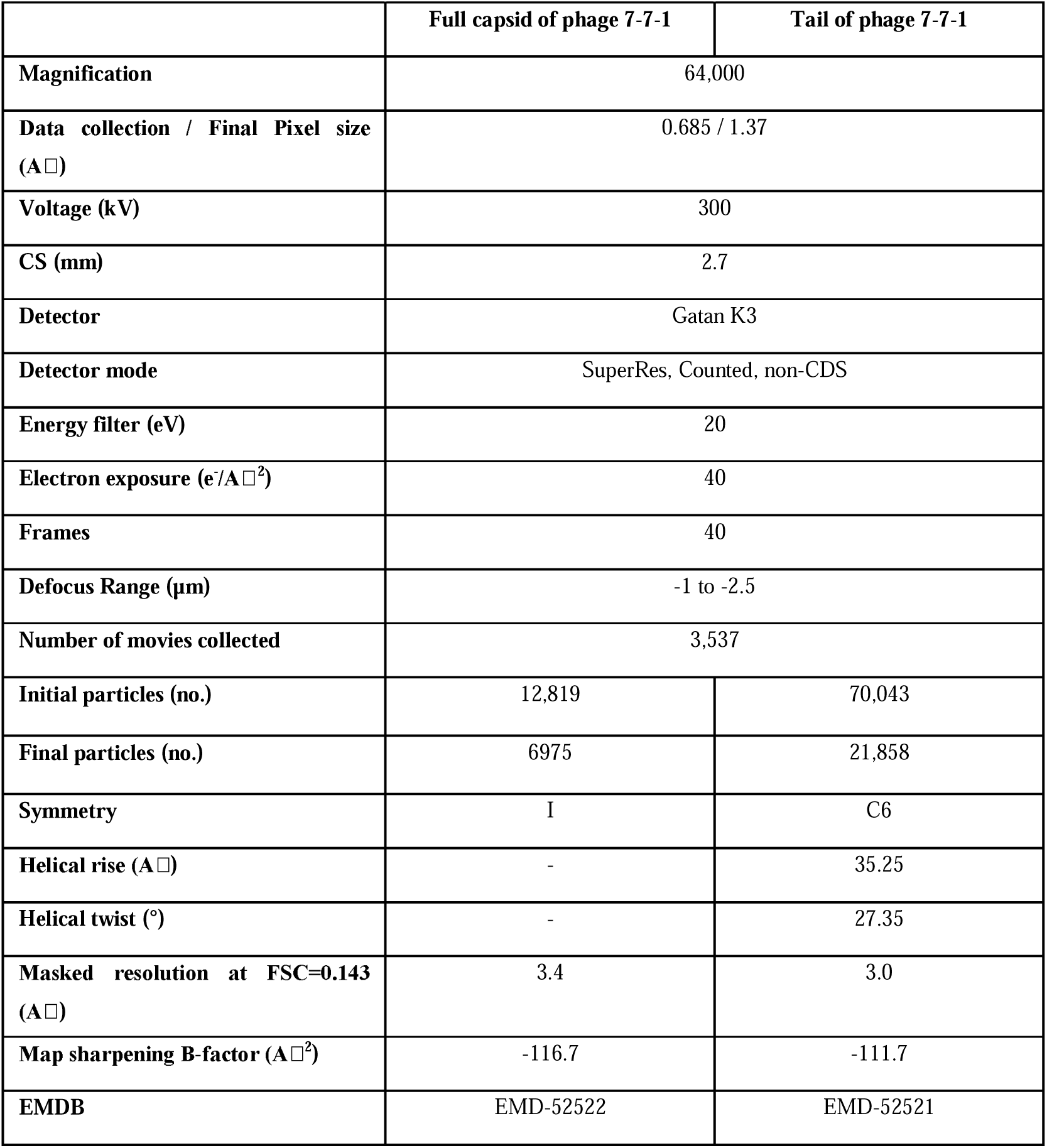
Cryo-EM data collection parameters and processing.

**Supplementary Table. 2.** Model refinement and validation statistics.

|  | **Pentamer-hexamers**<br>(1 pentamer, 1 neighbor<br>hexamer, 1 further<br>neighbor hexamer, with<br>decoration proteins and<br>linking proteins) | **Tail**<br>(2 layers, 12 sheath proteins and<br>12 inner tube proteins) |
| --- | --- | --- |
| **PDB** | 9HZ8 | 9HZ7 |
| **Model compositions** |  |  |
| Amino Acids | 9694 | 7596 |
| **R.m.s. deviations** |  |  |
| Bond lengths (Å) | 0.003 | 0.003 |
| Bond angles (°) | 0.539 | 0.569 |
| **Validation** |  |  |
| MolProbity score | 1.33 | 1.43 |
| Clash score | 3.37 | 3.15 |
| Rotamers outliers (%) | 0.00 | 0.00 |
| **Ramachandran plot** |  |  |
| Favored (%) | 96.72 | 95.4 |
| Allowed (%) | 3.27 | 4.6 |
| Outliers (%) | 0.01 | 0.00 |
| **Rama-Z** |  |  |
| Whole | -0.56 (0.08) | -0.49 (0.10) |
| Helix | 1.18 (0.18) | 0.24 (0.14) |
| Sheet | 0.21 (0.10) | 0.33 (0.12) |
| Loop | -0.95 (0.08) | -0.81 (0.09) |
| **Model vs. Data** |  |  |
| CC (mask) | 0.84 | 0.85 |
| CC (volume) | 0.82 | 0.78 |

**Supplementary Figure 1.**
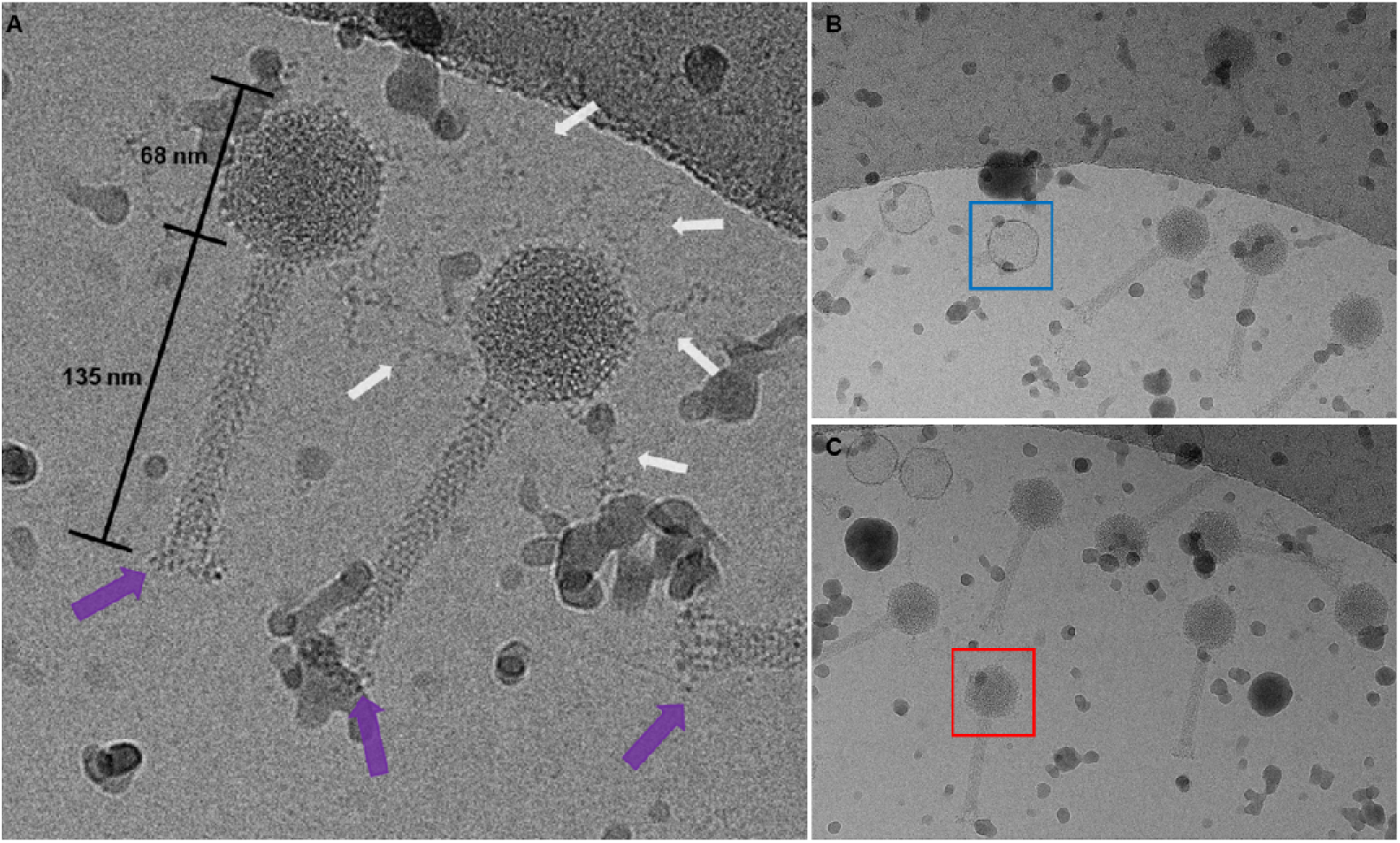
Representative micrographs of phage 7-7-1. **(A)** An enlarged image of a representative cryo-EM image from SPA. This image showcases a hexagonal head with a diameter around 68 nm and a contractile tail measuring 135 nm in length. Notably, the end of the contractile tail is decorated with abundant bushy tail fibers (purple arrows), and many long-curved capsid fibers surrounding the capsid (white arrows). **(B)** A representative micrograph of phage 7-7-1 with empty capsids (blue rectangle). **(C)** A representative micrograph of phage 7-7-1 with full capsids containing DNA (red rectangle). Notably, most of the phages observed in our dataset were intact capsids filled with DNA.

**Supplementary Figure 2.**
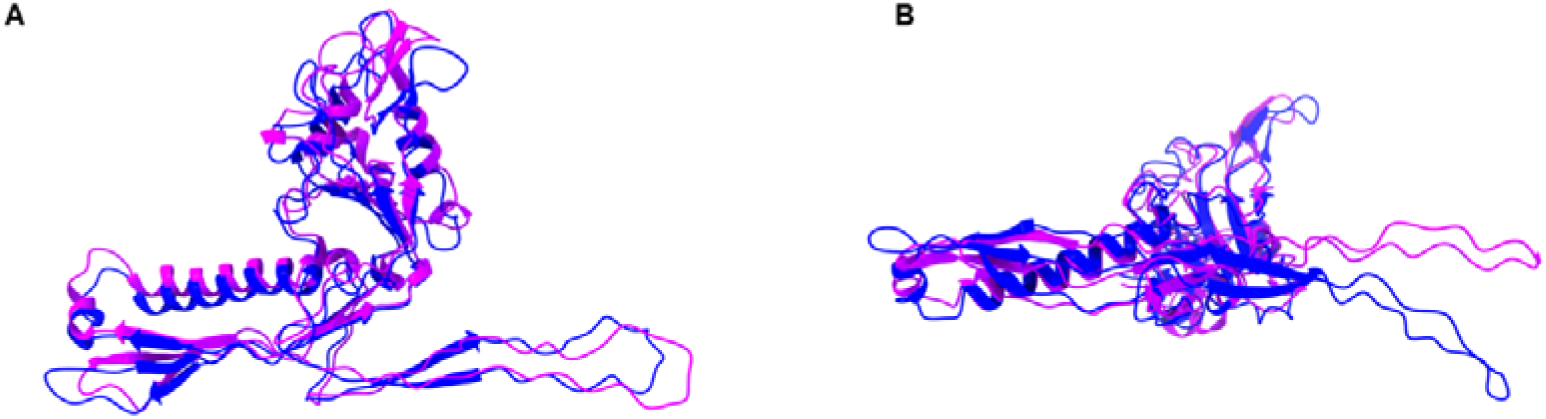
Structural comparison between the MCP (purple) in a hexamer and MCP (dark blue) in a pentamer. There are only small differences, but the MCP (dark blue) in a pentamer is more curved than the MCP in a hexamer. The right image in **(B)** shows the comparison after rotation of the left structure by 90°.

**Supplementary Figure 3.**
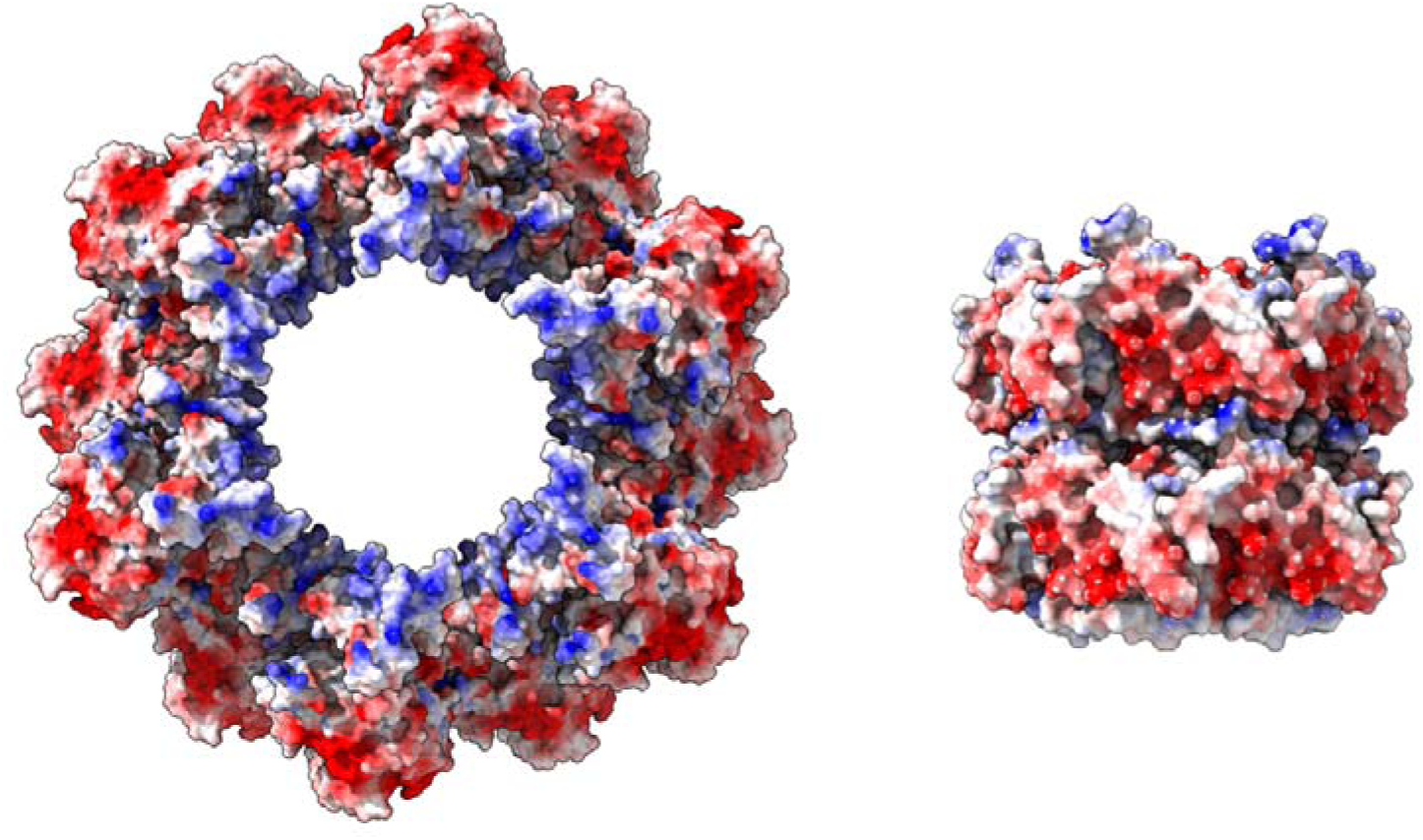
Electrostatic interactions between phage sheath and tail. The tail sheath has a negatively charged interior (left), which interacts with the positively charged exterior of the tail tube (right).

**Supplementary Figure 4.**
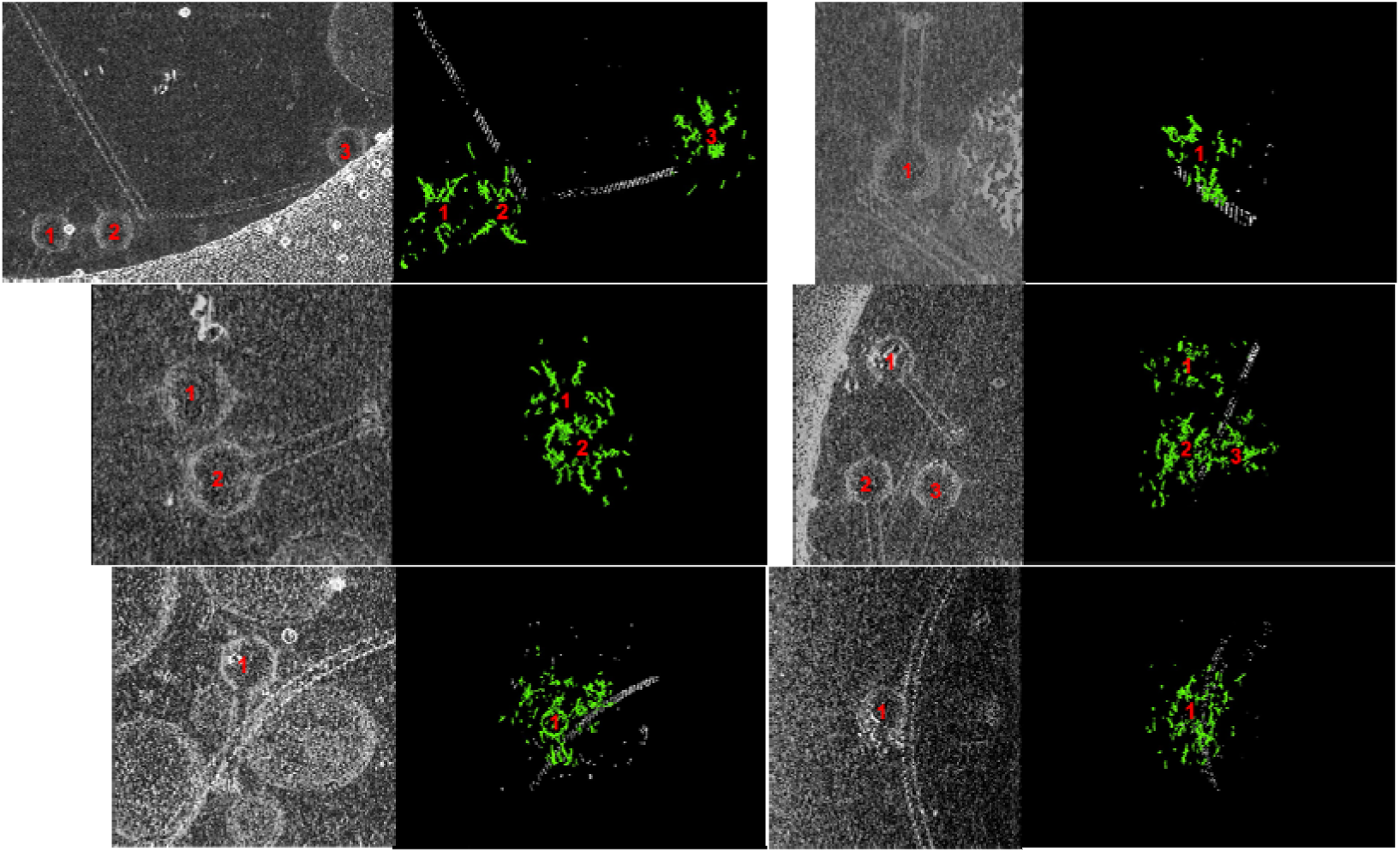
Trimmed tomograms of phage 7-7-1 attached to flagella and 3D structures of capsid fibers. Six groups of trimmed tomograms and their corresponding 3D structures. In each group, the left figure displays a trimmed tomogram, which is overlaid and visualized into 2D image, created through the 3D projection function in FIJI. On the right, the 3D structures of capsid fibers (green) and flagella (gray) are visualized, generated using the neural network. Red numbers are marked at the center of phage capsids for location and identification purposes.

**Supplementary Figure 5.**
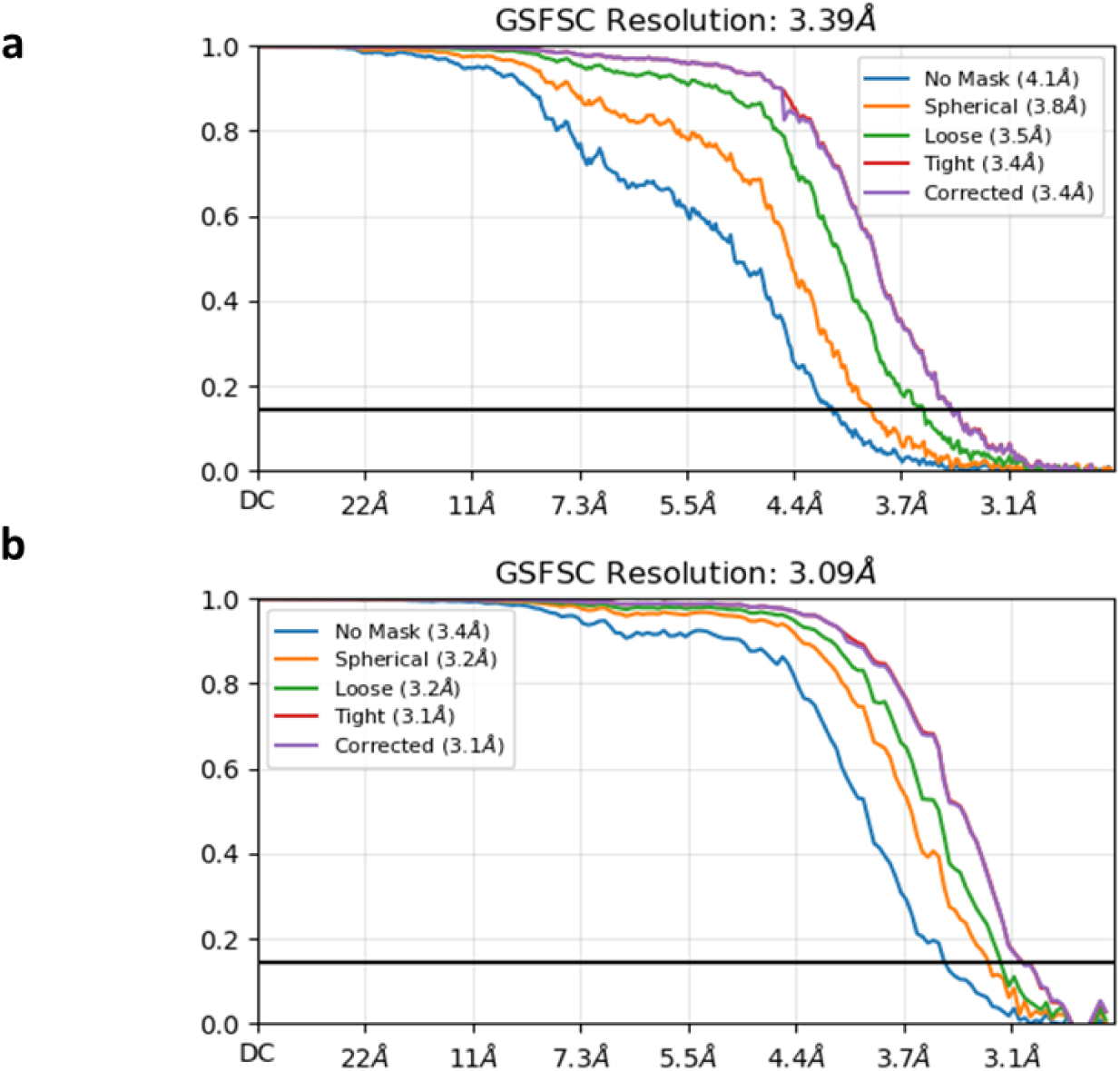
Fourier shell correlation curves. Map-map FSC curves of the phage 7-7-1 capsid (a) and tail (b) at a cut-off of 0.143.

## Notes

### Competing Interest Statement

The authors have declared no competing interest.

